# Relieving platelet inhibition using a novel bi-specific antibody: A novel approach for circumventing the platelet storage lesion

**DOI:** 10.1101/2025.03.20.644378

**Authors:** Alyssa J. Moroi, Peter J. Newman

**Affiliations:** Blood Research Institute Versiti, Milwaukee, WI; Department of Pharmacology & Toxicology Medical College of Wisconsin, Milwaukee, WI; Department of Cell biology, Neurobiology, and Anatomy Medical College of Wisconsin, Milwaukee, WI

**Author notes:** **Address correspondence to:** Peter J. Newman, Ph.D., Versiti Blood Research Institute, Versiti Blood Center of Wisconsin, 8727 W. Watertown Plank Rd, Milwaukee, WI 53226, **Email:**.

**Keywords:** Platelets, GPVI, Platelet activation, PECAM-1, CD148

## Abstract

**Background:** Human platelets experience structural and functional deterioration during extracorporeal storage at either room temperature or in the cold, impairing their reactivity and diminishing their hemostatic effectiveness following transfusion. PECAM-1 is an inhibitory receptor on platelets that exerts its inhibitory effects via phosphorylation of tyrosine residues that lie within its cytoplasmic immunoreceptor tyrosine-based inhibitory motifs (ITIMs). The purpose of this investigation was to attempt to restore platelet reactivity by impairing the inhibitory activity of PECAM-1.

**Methods:** To counteract PECAM-1-mediated inhibition, we developed a novel bispecific tandem single-chain variable fragment (scFv) that ligates the protein-tyrosine phosphatase, CD148, with PECAM-1, promoting dephosphorylation of PECAM-1 ITIMs. We then analyzed the ability of this engineered tandem scFv (taFv 179) to improve adhesion and aggregation responses in vitro and under conditions of flow.

**Results:** Addition of taFv 179 enhanced secretion, aggregation, and activation responses of both freshly isolated and stored platelets, particularly in response to weak agonists. taFv 179 also improved thrombus formation on collagen-coated surfaces under conditions of arterial flow.

**Conclusions:** These findings demonstrate that enforced approximation of a phosphatase next to PECAM-1 ITIM tyrosines receptors is a novel strategy for enhancing the functionality of stored platelets, with potential implications for improving the effectiveness of platelet transfusions.

**Essentials:**

- Platelets lose reactivity upon extracorporeal platelet storage.
- Platelets lacking PECAM-1 are known to be hyperresponsive to platelet agonists.
- Addition of a bispecific antibody that phenocopies PECAM-1 deficiency partially restores the reactivity of stored platelets.

## Introduction

Platelets play a central role in thrombosis and hemostasis. Platelet transfusion is a common therapeutic approach for patients with thrombocytopenia, which may result from malignancy, chemotherapy, blood loss due to trauma or surgery, infections, or inherited platelet disorders. Currently, platelets collected for transfusion are stored at room temperature (20–24°C) with contin-uous agitation. However, their shelf-life is limited to five days due to bacterial contamination and the development of the so-called platelet storage lesion (PSL), which impairs platelet function. The PSL is characterized by altered metabolism, increased apoptosis, morphological changes, and reduced membrane integrity, together leading to impaired aggregation, reduced clot stability, and shortened circulation lifespan upon transfusion. These changes limit the efficacy of stored platelets in controlling bleeding.^1-5^

There have been numerous attempts to extend the duration of platelet storage, with refrigerated platelets having both the longest history^6^ and having the most promise in military and trauma settings, both of which require readily available blood products in remote or austere environ-ments^7,8^ Cold-stored platelets can be kept at 1–6°C for 7–14 days, but still exhibit marked functional defects, including clustering, desialylation, and loss of surface membrane glycoproteins, leading to reduced circulation time following transfusion^9^ thereby limiting their clinical application.^5,10,11^ In an effort to restore cellular activation, bispecific diabodies capable of targeting and inactivating inhibitory receptors have recently been developed for other cell types. Such reagents work by recruiting a transmembrane phosphatase next to a receptor tyrosine kinases, effectively inhibiting both tonic and ligand-activated inhibitory signaling.^12,13^ For example, a bispecific diabody targeting programmed cell death-1 (PD-1) and CD45 represses PD-1 signaling in T cells, imparting improved therapeutic efficacy in preclinical cancer models.^12^

PECAM-1 is a member of the Ig inhibitory receptor family^14,15^ whose inhibitory function is mediated by two cytoplasmic domain immunoreceptor tyrosine-based inhibitory <u>m</u>otifs (ITIMs) that become phosphorylated upon cellular activation, leading to recruitment of downstream signal-ing molecules like SHP-1 and SHP-2^16,17^ These phosphorylation-dependent PECAM-1 signaling complexes function to *negatively* regulate platelet activation, as evidenced by studies both in vitro,^18^ and in PECAM-1-deficient mice, which demonstrate *hyper*-responsiveness to GPVI agonists, enhanced thrombus formation, and increased thrombus stability.^19-21^ CD148, on the other hand is a receptor-type tyrosine phosphatase (RPTP) that acts as a *positive* regulator of platelet activation, and CD148-deficient mice exhibit *impaired* platelet activation and thrombus forma-tion.^22,23^

Given their opposing effects on platelet reactivity, and the function of CD148 as a transmembrane tyrosine phosphatase, we hypothesized that bringing CD148 into close approximation with PECAM-1 might maintain PECAM-1 cytoplasmic ITIMs in a dephosphorylated state, thereby phenocopying PECAM-1 deficiency, potentiating platelet activation, and partially restoring some of the function lost upon platelet storage. We therefore engineered a novel bispecific tandem single-chain variable fragment (taFv) that binds both PECAM-1 and CD148 that is able to enhance activation responses in both freshly isolated and stored platelets, and demonstrate that it does so by reducing PECAM-1 ITIM phosphorylation. Taken together, these results provide a novel strategy for improving platelet functionality during storage and transfusion.

## Methods

### Reagents

FITC-conjugated PAC-1, and PE-conjugated P-selectin were from BD Biosciences (San Jose, California, United States). Antibody against human CD148 was from R&D systems (Minneapolis, Minnesota, United States). Antibody against human PECAM-1 (PECAM-1.3)^24^ was produced by the Versiti Blood Research Institute Core Labs. Antibodies against Src-pTyr_527_, Lyn-pTyr_507_, and ERK-pThr_202_/Tyr_204_ were from Cell Signaling Technology (Danvers, Maryland, United States). Antibody against human GPVI was from Blood Research Institute Hybridoma Core Laboratory. Antibody against actin was from Thermo Fisher Scientific (Waltham, Massachusetts, United States). Horseradish peroxidase-conjugated secondary antibodies were from Jackson Immuno Research (West Grove, Pennsylvania, United States). Collagen-related peptide (CRP) and PAR-1 agonist, thrombin receptor-activating peptide (TRAP) was prepared by the Blood Research Institute Protein Core Laboratory. Collagen was from Chrono-Log corporation (Havertown, Pennsylvania, United States). All other reagents were from Sigma-Aldrich (St Louis, Missouri, United States).

### Construction of bispecific tandem scFv

A DNA fragment with the light- and heavy-chain variable sequences (VL and VH) of CD148 mAb (Ab-2 from patent US7449555B2) were recombinantly joined to the VL and VH of anti-PECAM1 mAb PECAM1.3 by a linker consisting of (Gly-Gly-Gly-Gly-Ser)_3_ and synthesized (Twist Bioscience). C-terminal histidine (His) tag was added to facilitate protein purification. The resulting bispecific tandem scFv construct is in following arrangement: signal peptide-CD148 VL-(G4S)_3_-CD148 VH-SGGGGS-PECAM-1 VL-(G4S)_3_-PECAM-1 VH-GGG-8×His.

The DNA fragment was inserted to pcDNA3.4 (Invitrogen, Waltham, Massachusetts, United States) according to the manufacturer’s protocol.

### Cell culture and expression of bispecific tandem scFv

Expi293F cells were cultured in Expi293 expression medium (Gibco, Waltham, Massachusetts, United States) at 37°C, 8%CO_2_. The expression vector was transfected into Expi293F cells using FectoPro reagent (Polyplus, New York, New York, USA). Briefly, the mixture of 80 µg DNA and 80 µL of FectoPro was used to transfect 100 mL of 3–4×10^6^ cells/mL. And cells were grown for 4 days at 37°C on a shaker. The protein was purified from culture supernatant by affinity chromatography using Ni-NTA Sepharose beads (Thermo Fisher Scientific) according to the manufacturer’s protocol. After purification, purified fractions were concentrated and buffer exchanged to PBS. Protein purity was determined by SDS-PAGE.

### Platelet isolation

All donors gave informed consent, and the study was approved by the Medical College of Wisconsin Institutional Review Board. Blood was collected from healthy donors into anticoagulant sodium citrate. Platelet-rich plasma (PRP) was prepared by centrifugation. Washed platelets were prepared as previously described^25^ with modification in Tyrode’s buffer (137 mM sodium chloride, 2.5 mM potassium chloride, 0.36 mM of sodium dihydrogen phosphate dihydrate, 13.8 mM sodium bicarbonate, 20 mM N-2-hydroxyethylpiperazine-N9-2-ethanesulfonic acid, and 0.1% glucose). The pellet of washed platelets was resuspended in Tyrode’s buffer to an appropriate concentration (2×10^8^ mL^-1^) for the experiments. For room temperature storage, PRP and washed platelets were stored with gentle agitation, and cold stored platelets were stored without agitation for indicated days.

### Platelet aggregation and secretion

Aggregation was monitored by light transmission with a Born Lumi-Aggregometer (Chrono-log Corporation).

### Flow cytometry

Human platelet integrin αIIbβ3 activation and α-granule secretion were measured by flow cytometry on a BD Accuri™ C6 plus (BD Biosciences) flow cytometer. The FITC-conjugated PAC-1 antibody, which binds the activated integrin αIIbβ3 and PE-conjugated antibody binds the α-granule marker P-Selectin were used. Platelets (5×10^5^) were stimulated as indicated and fixed using 2% paraformaldehyde, then read on the flow cytometer until a total of 10,000 events were collected from each well.

### In vitro platelet adhesion and thrombus formation in microfluidic chambers

Whole blood was collected and anticoagulated with heparin/PPACK. PRP was isolated by centrifugation treated with taFv 179 (20 µg/mL) 37ºC, 30 min and labeled with mepacrine (Calbiochem, San Diego, California, United States). Red blood cells were collected after centrifugation and washed with PBS. Before the experiment red blood cells (4×10^6^ mL^-1^) were added to the PRP. Thrombus formation on collagen was performed as previously described on collagen (50 µg/mL) at an arterial sheer rate of 1500 s^-1^. Images of platelet adhesion and thrombus formation were acquired by epifluorescence microscopy at a rate of 1 frame per second. Platelet accumulation was analyzed using IMAGE J software (NIH, Bethesda, Maryland, United States), and results are reported as the mean area covered by platelet aggregates (%). Area under curve (AUC) was analyzed using Prism 10.1.0 software (GraphPad, San Diego, California, United States).

### Western blotting

Washed platelets were stimulated with collagen-related peptide (CRP; 5 µg/mL) at 37°C with stirring for the indicated times, and reactions were terminated by adding an equal volume of ice-cold 2×lysis buffer (300 mM of sodium chloride, 20 mM of tris[hydroxymethyl]aminomethane, 2 mM EGTA, 2 mM EDTA, and 2% NP40 [pH, 7.5]). The samples were diluted with 4×Laemmi sample buffer (BIO-RAD) separated by SDS-PAGE (sodium dodecyl-sulfate polyacrylamide gel electrophoresis), and transferred to a polyvinylidene difluoride membrane. Western blotting was performed with the indicated antibodies. Bands were quantitated by densitometry using IMAGE J software.

### Statistics

Statistical analyses were performed with the two-way ANOVA or paired-t test as indicated using Prism 10.1.0 software. All data expressed as the mean±standard error of the mean (SEM).

## Results

### Development of bispecific tandem scFv (taFv 179)

The receptor inhibition by phosphatase recruitment (RIPR) approach has previously been reported as a strategy to potentiate T cell activation.^12^ This mechanism involves engaging phosphatases to modulate receptor signaling, presenting a novel avenue for therapeutic development. To investigate whether this approach could be adapted to enhance platelet activation, we developed a bispecific taFv targeting key platelet receptors. The bispecific construct, termed taFv 179, was designed to bind both the platelet ITIM receptor PECAM-1 and the RPTP CD148. By simultaneously targeting these receptors, taFv 179 aims to enhance platelet activation through the RIPR mechanism. The variable regions of PECAM-1 monoclonal antibody (PECAM1.3) and CD148 monoclonal antibody (Ab-2; Amgen patent) were utilized to generate the tandem scFv construct. The resulting taFv 179 molecule was engineered with a C-terminal 8×His tag to facilitate both purification and detection during downstream analysis (**Figures 1A and 1B**). Purification of taFv 179 yielded a protein with a molecular weight of 53 kDa, as determined by SDS-PAGE (**Figure 1C**). To evaluate its functional binding capability, taFv 179 was tested for its ability to bind to human platelets. Flow cytometry analysis demonstrated strong binding of taFv 179 to human platelets, validating its bispecific targeting properties (**Figure 1D**).

**Figure 1.**
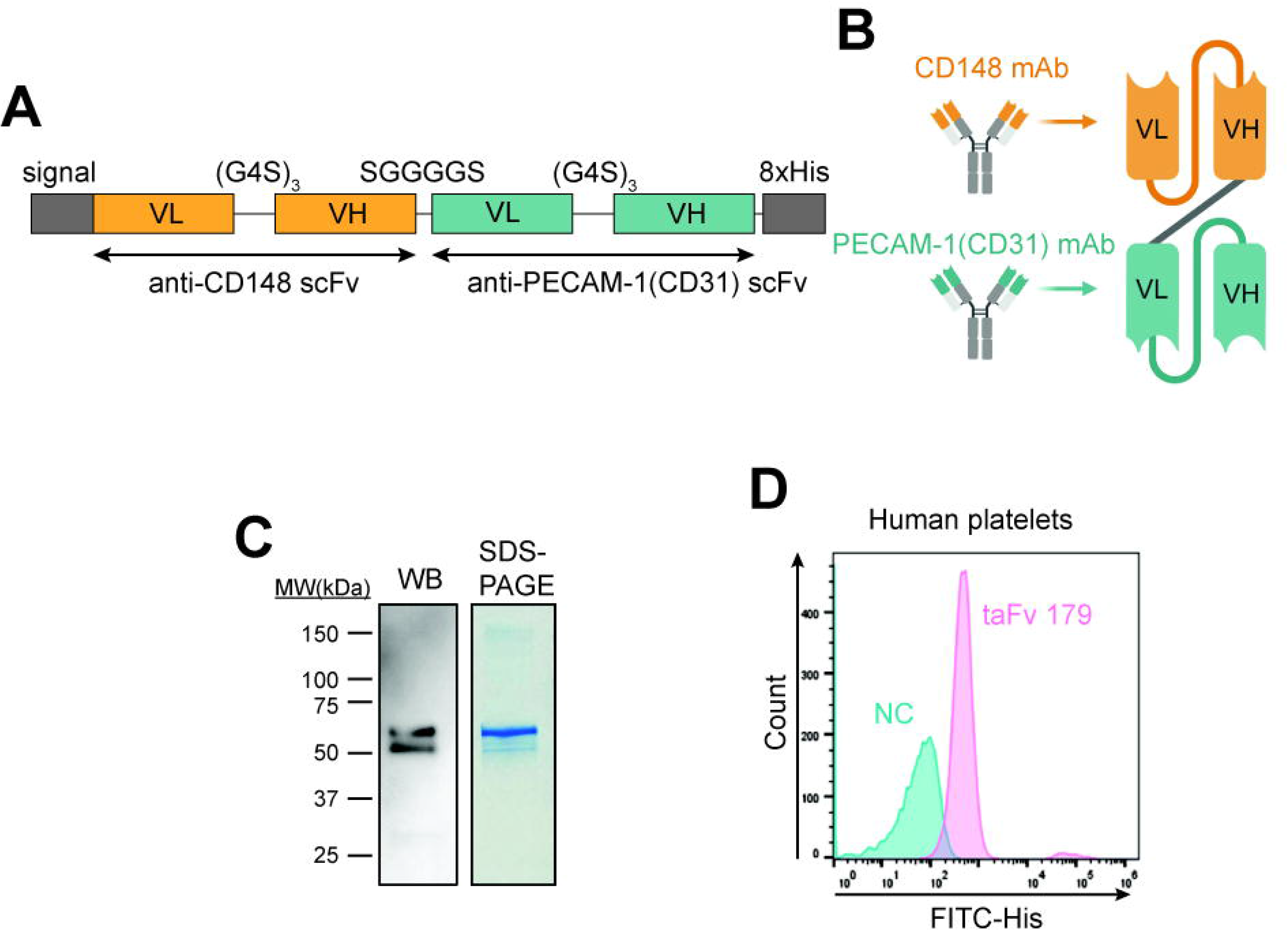
Design and production of the tandem single-chain antibody (taFv) 179. (**A**) Arrangement of gene modules in the pcDNA3.4 vector. Signal denotes the leader sequence from the human immunoglobulin heavy chain. The C-terminus of taFv contains a His tag. (**B**) taFv 179 is a bispecific tandem scFv composed of an N-terminal CD148 antibody-targeting scFv connected by a short flexible linker to a PECAM-1 antibody-binding scFv. Both scFvs are defined as linear assemblies of variable heavy (VH) and variable light (VL) chains. (**C**) Recombinant taFv 179 was produced in Expi293F cells and purified by Ni-NTA affinity chromatography. Its purity was confirmed by SDS-PAGE. The final mature protein is 53.62 kDa. (**D**) Binding of taFv 179 to human platelets compared to a secondary antibody only control (NC).

### taFv 179 potentiates platelet activation

PECAM-1 is an ITIM-bearing receptor that acts predominantly to inhibit the function of ITAM-bearing activating receptors like the GPVI/FcRγ-chain complex,^19,26^ though lesser effects on other agonists have been found^18,20^ To determine whether taFv 179 could potentiate platelet activation, washed human platelets were pretreated with various concentrations of taFv 179 before activation with the GPVI agonists collagen-related peptide (CRP) and collagen, and the GPCR agonist, thrombin or TRAP. taFv 179 treatment itself did not activate platelets (not shown), however, as shown in **Figures 2A and 2B**, taFv 179 enhanced platelet aggregation in a dose-dependent manner. We further assessed platelet activation by measuring P-selectin exposure and PAC-1 binding (indicative of active integrin αIIbβ3) following stimulation. taFv 179 treatment significantly enhanced platelet activation, as demonstrated by an increased P-selectin/PAC-1-positive population and elevated PAC-1 mean fluorescence intensity (MFI) after CRP stimulation in both washed platelets (**Figure 2C**) as well as platelets in PRP (**Figure 2D**).

**Figure 2.**
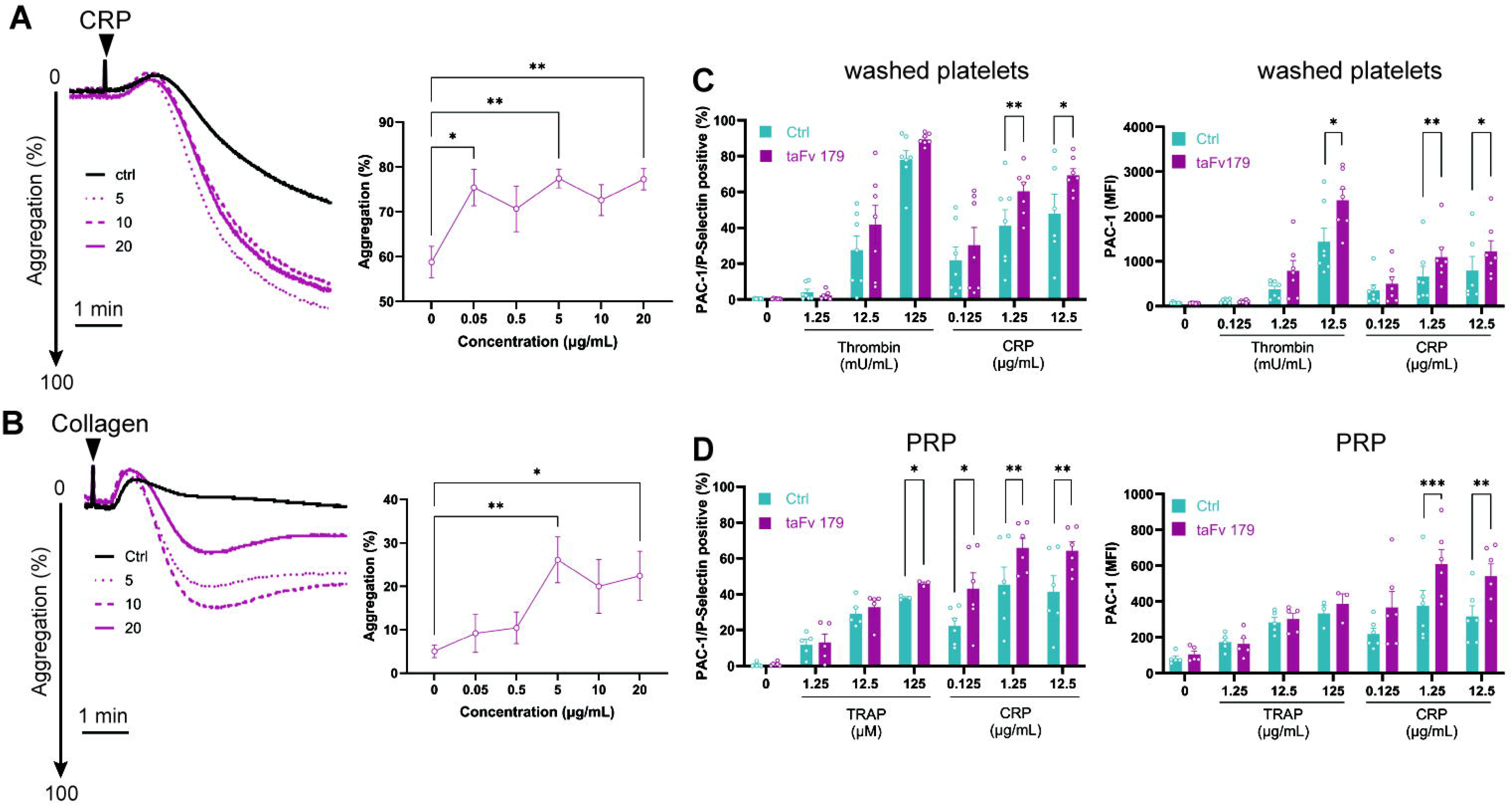
taFv 179 enhances GPVI-mediated platelet aggregation. Freshly isolated washed human platelets were treated with the indicated concentrations of taFv 179 (µg/mL) for 20 min at 37 °C. Aggregation was assessed by lumiaggregometry in response to (**A**) CRP (5 µg/mL) or (**B**) collagen (0.5 µg/mL). Representative traces and maximum aggregation (%) are shown (N=6–11). Statistical analysis was performed using two-way ANOVA (*P < 0.05, **P < 0.01 compared to control [Ctrl]). Freshly isolated washed human platelets (**C**) or platelet-rich plasma (PRP, (**D**)) were treated with taFv 179 (20µg/mL) for 20 min at 37 °C. P-selectin exposure and PAC-1 binding (indicators of integrin αIIbβ3 activation) were measured by flow cytometry after stimulation with thrombin, TRAP, or CRP at the indicated concentrations. Data represent the mean ± SEM of PAC-1/P-selectin-positive populations and PAC-1 mean fluorescence intensity (MFI) (N = 3–7). Statistical analysis was performed using a paired t-test (*P < 0.05, **P < 0.01, ***P<0.001 compared to Ctrl).

Platelet storage is known to impair activation, which presents challenging limitations in transfusion settings. Therefore, we investigated whether taFv 179 could restore some of the functionality of stored platelets. As shown in **Figure 3**, addition of taFv 179 had a modest, but statistically significant, enhancement on the ability of platelets stored at room temperature for either 2 or 5 days to aggregate in response to CRP – the agonist most likely to be inhibited by PECAM-1.^19^ Potentiation was seen whether the platelets had been stored in buffer or in PRP. These results were corroborated by increased exposure of P-selectin and activated GPIIb-IIIa on the platelet surface in the presence of taFv 179 (**Figure 4**). This potentiation was primarily seen in platelets stored at room temperature, with more modest improvement for platelets stored in the cold.

**Figure 3.**
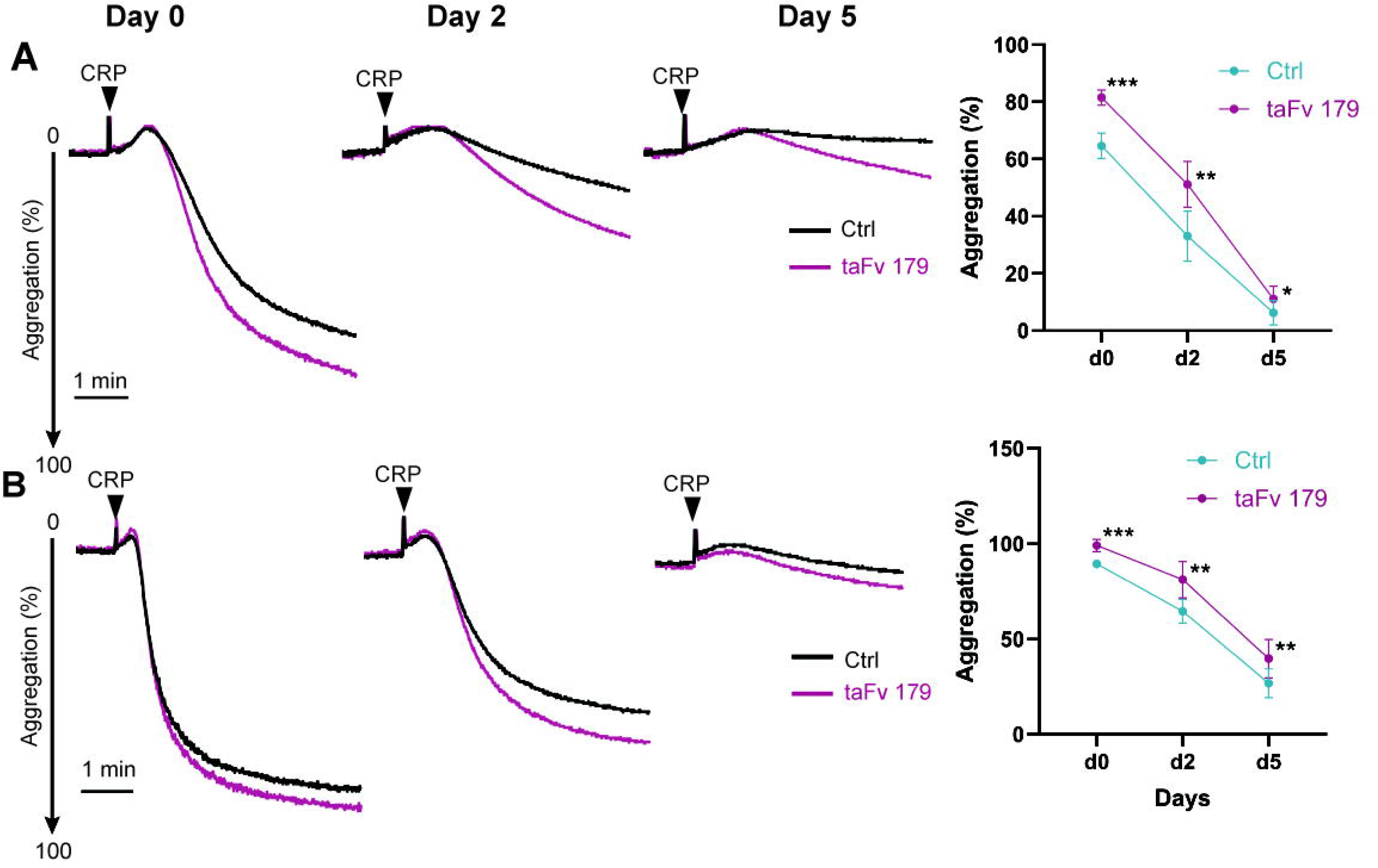
taFv 179 treatment potentiates GPVI-mediated platelet aggregation. (**A**) Freshly isolated washed platelets and (**B**) platelet-rich plasma (PRP) stored at room temperature for 2 or 5 days were treated with taFv 179 (20 µg/mL) for 20 min at 37 °C. Aggregation was assessed by lumiaggregometry in response to CRP (5 µg/mL). Representative traces and maximum aggregation (%) are shown (N = 5–12). Statistical analysis was performed using a paired t-test (*P < 0.05, **P < 0.01, ***P<0.001 compared to control [Ctrl])

**Figure 4.**
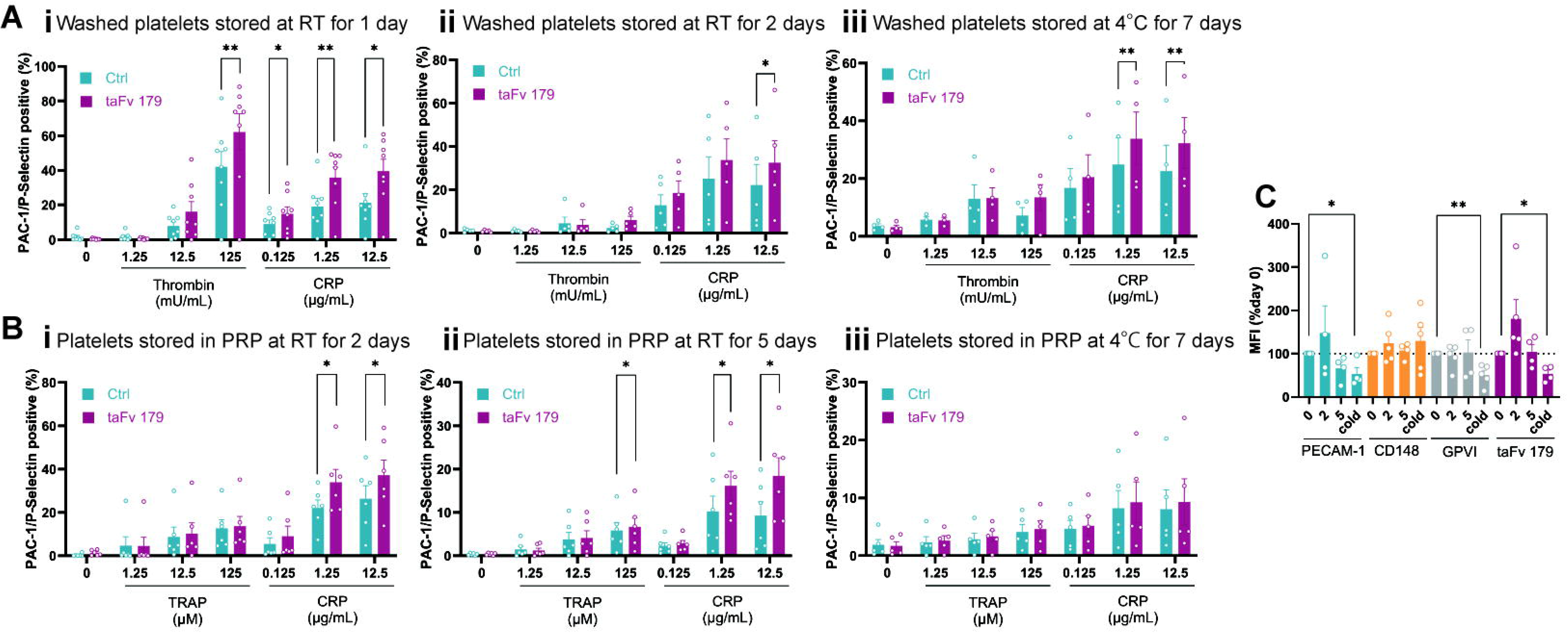
taFv 179 potentiates activation of stored human platelets. Washed platelets (**Panels A i, ii and iii**) and platelets in PRP (**Panels B i, ii and iii**) were stored at room temperature for 1 (**panel A i**), 2 (**panels A ii and B i**) or 5 (**panel B ii**) days at room temperature, or for 7 days at 4 °C (**panels iii**). All samples were then incubated with vehicle or taFv 179 (20 µg/mL) for 20 min at 37 °C. Following stimulation with thrombin, TRAP, or CRP at the indicated concentrations, P-selectin exposure and PAC-1 binding (indicators of integrin αIIbβ3 activation) were measured by flow cytometry. Data represent the mean ± SEM of PAC-1/P-selectin-positive populations (N = 4–8). Statistical analysis was performed using a paired t-test (*P < 0.05, **P < 0.01 compared to control [Ctrl]). (**C**) PRP was stored at room temperature for 0, 2, and 5 days, or at 4 °C for 7 days. Binding of mAbs to PECAM-1, CD148, GPVI, as well as taFv 179 are depicted as a % of binding at Day 0. (n = 4–5). Statistical analysis was performed using a paired t-test (*P < 0.05,**P<0.01 compared to day 0)

Cold storage of platelets has been shown to lead to a decrease in the surface expression of GPVI, impairing responses to its ligands, collagen and GPVI.^10^ To further investigate the impact of storage on receptor expression, we measured surface expression levels of PECAM-1, CD148, and GPVI before and after room temperature and cold storage. As shown in **Figure 4C**, whereas PECAM-1, CD148, and GPVI levels remained stable during room temperature storage, both GPVI and PECAM-1 decreased during 7 days of cold storage, as did the ability of taFv 179 to bind. These findings are consistent with the relatively modest improvement in platelet reactivity afforded by taFv 179 to enhance the reactivity of cold-versus room temperature-stored platelets (compare panel ii with panel iii of **Figure 4B**).

### taFv 179 potentiates thrombus formation on collagen

To examine potential of taFv 179 to improve hemostasis under conditions of flow, freshly prepared PRP, supplemented with red blood cells^27^, was subjected to arterial shear conditions in a microfluidic flow chamber. As shown in **Figure 5**, addition of taFv 179 significantly enhanced thrombus formation on a collagen-coated surface at an arterial shear rate of 1500 sL^1^. taFv 179 also improved the hemostatic effectiveness of PRP that had been stored at room temperature for 2 and 5 days (**Figures 6A and 6B**), but failed to improve thrombus formation of platelets that had been stored in the cold for 7 days (**Figure 6C**), consistent with the reduction of PECAM-1 surface expression and taFv 179 binding seen during cold storage (**Figure 4C**). Collectively, these findings demonstrate that taFv 179 is able to enhance thrombus formation on collagen under flow of both fresh and room temperature-stored platelets.

**Figure 5.**
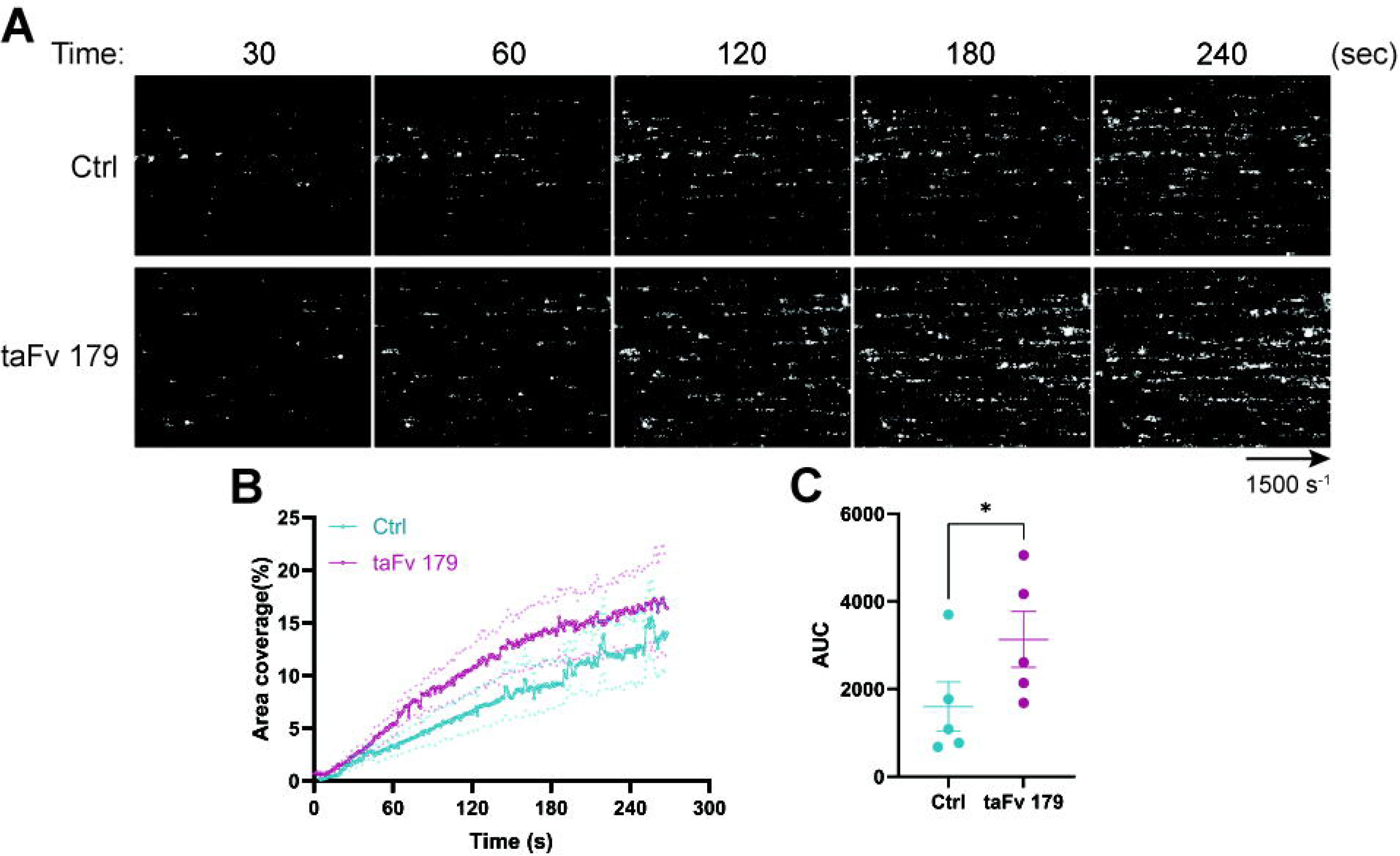
taFv 179 improves platelet adhesion to collagen under conditions of flow. (**A**) Platelet-rich plasma (PRP) was treated with vehicle or taFv 179 (20 µg/mL) for 30 minutes at 37 °C. PRP was labeled with mepacrine, mixed with red blood cells (4 × 10L µL^-1^), and perfused over collagen-coated microchannels under arterial shear (1500 sL^1^). Images were captured using a 20× objective. (**B**) Surface area coverage and (**C**) area under the curve (AUC) were quantified and reported as mean ± SEM (N = 5). Statistical analysis was performed using a paired t-test (*P<0.05 compared to control [Ctrl]).

**Figure 6.**
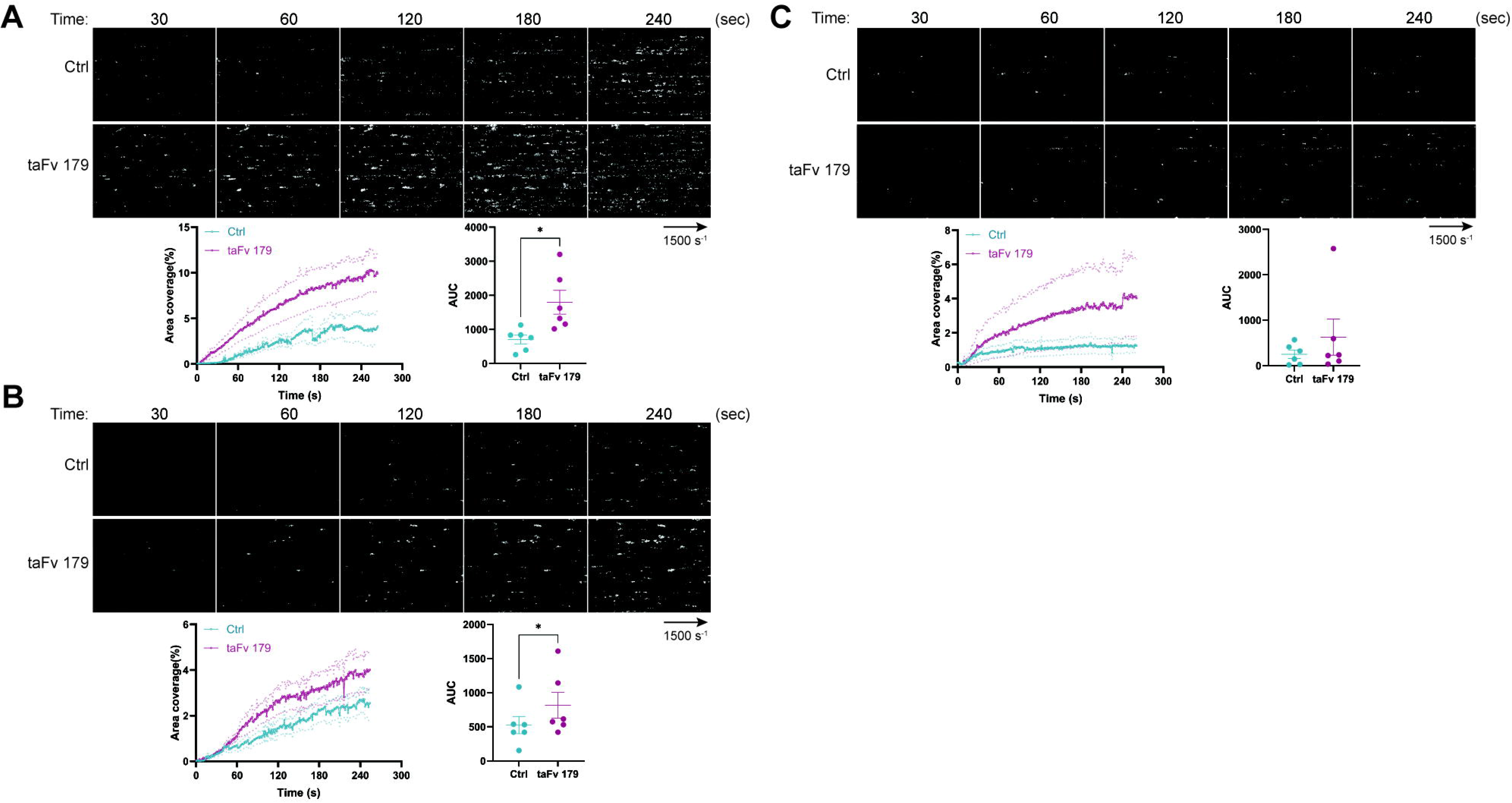
Increased hemostatic function of stored platelets in the presence of taFv 179. Platelet-rich plasma (PRP) stored at room temperature for (**A**) 2 days, (**B**) 5 days, or at (**C**) 4°C for 7 days was treated with vehicle or taFv 179 (20 µg/mL) for 30 minutes at 37 °C. PRP was labeled with mepacrine, mixed with red blood cells (4 × 10L µL^-1^), and perfused over collagen-coated microchannels under arterial shear (1500 sL^1^). Flow chamber images were captured using a 20× objective. A single representative flow chamber image and its densitometric quantitative curve is shown for each condition. Surface area coverage and area under the curve (AUC) were quantified and reported as mean ± SEM (n = 6). Statistical analysis was performed using a paired t-test (*P < 0.05 compared to control [Ctrl]).

### taFv 179 enhances GPVI-signaling events by reducing inhibitory PECAM-1 ITIM phosphorylation

taFv 179 was designed to potentiate platelet activation by cross-linking CD148 and PECAM-1, thereby reducing PECAM-1 ITIM tyrosine phosphorylation. To investigate the changes in signaling events induced by taFv 179, platelets were pretreated with taFv 179 and then stimulated with CRP. As expected, taFv 179 significantly reduced PECAM-1 ITIM phosphorylation following GPVI stimulation (**Figures 7A and 7B**). Consistent with the reduction in PECAM-1 inhibitory signaling, there was a significant increase in GPVI-mediated signaling events, as reported by elevated phosphorylation of ERK. Interestingly, taFv 179 treatment also led to a decrease in C-terminal inhibitory tyrosine phosphorylation of Src (Tyr_527_) and Lyn (Tyr_507_), suggesting that these kinases are maintained in a pre-activated state in the presence of taFv 179. Taken together, these findings are consistent with the notion that taFv 179 is able to mechanistically phenocopy PECAM-1 deficiency by reducing PECAM-1 ITIM phosphorylation, thereby mitigating its negative regulatory effects on GPVI-mediated platelet activation.

**Figure 7.**
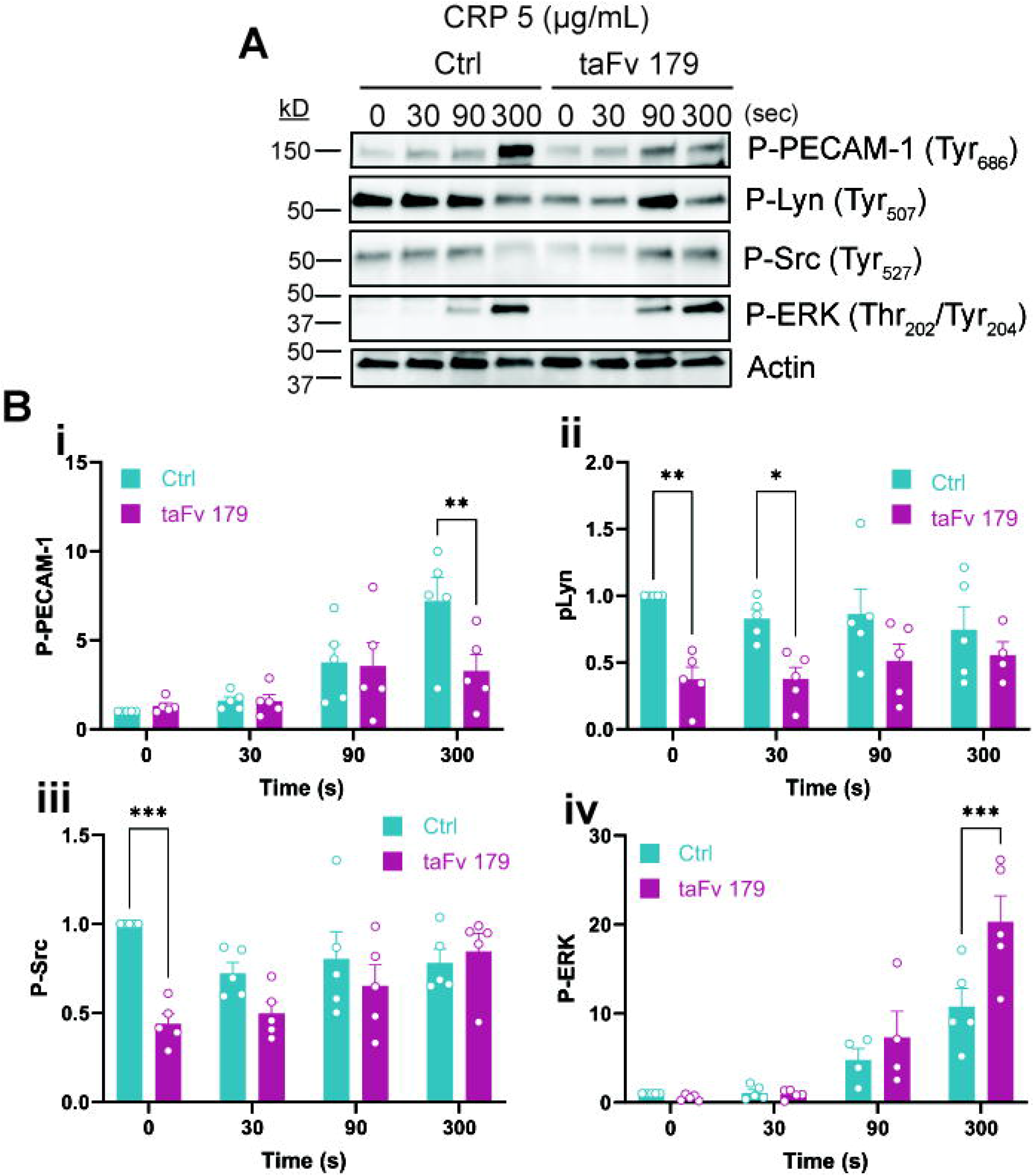
taFv 179 enhances GPVI-mediated signaling by dampening PECAM-1 inhibitory signaling. (**A**) Freshly isolated washed platelets were treated with vehicle or taFv 179 (20 µg/mL) for 20 minutes at 37 °C. Platelets were stimulated with 5 µg/mL collagen-related peptide (CRP) for 30, 90, and 300 seconds under stirring conditions in an aggregometer. Lysates were analyzed for phosphorylated PECAM-1 (P-PECAM-1; Tyr_686_), P-Lyn (Tyr_507_), P-Src (Tyr_527_), P-ERK (Thr_202_/Tyr_204_), and actin. The blot shown is representative of 4–5 independent experiments. (**B**) Band densities were quantified, normalized to actin, and further normalized to the control zero-second time point across all experiments. Data are presented as mean ± SEM (n = 4–5). Statistical analysis was performed using the paired t-test (*P < 0.05, **P < 0.01, ***P<0.001 compared to control [Ctrl]).

## Discussion

In this study, we report the development and characterization of a bispecific antibody that targets the ITIM receptor PECAM-1 and the phosphatase CD148. This proof-of-concept report demonstrates that taFv 179 and proteins with similar binding properties are able to enhance the activation of both freshly isolated and room temperature-stored platelets. To our knowledge, this is the first study to generate a small protein capable of potentiating platelet activity, particularly in stored platelets. taFv 179, therefore, has the potential to improve the hemostatic effectiveness of platelets used in transfusions.

Bispecific scFvs offer several advantages over intact bispecific antibodies, including a small molecular size that reduces the likelihood of steric hindrance, and the absence of an Fc region to prevent binding and activation of cell surface Fcγ receptors. The absence of an Fc region, however, can render scFvs less stable, with a correspondingly shorter circulating half-life. Bispecific taFvs can be modified to contain a modified Fc region to improve stability. Other future modifications of taFv 179 could include developing a triabody version to compensate for the imbalance in the stoichiometry of PECAM-1 (copy number ∼9,500 receptors/platelets) and CD148 (copy number ∼2,800/platelet).^22,28^ The ultimate goal is to develop 2^nd^ and 3^rd^ generation products to be used in platelet transfusions for patients recovering from surgery or trauma, where enhanced platelet activation is needed, and to improve the utility of stored platelets by restoring their activity.

By bringing the transmembrane protein-tyrosine phosphatase, CD148, into close approximation PECAM-1, taFv 179 is able to reduce PECAM-1 ITIM phosphorylation following GPVI-mediated platelet activation, thereby phenocopying PECAM-1 deficiency, which has the effect of rendering platelets hyperresponsive to ITAM receptor-mediated activation responses.^15,19,21^ Interestingly, taFv 179 also reduced phosphorylation of the C-terminal inhibitory tyrosines of Src and Lyn, consistent with previous reports showing that CD148, along with the receptor-like protein tyrosine phosphatase CD45, dephosphorylates the inhibitory tyrosines of Lyn, thereby promoting B cell signaling.^29,30^ Moreover, studies in CD148-deficient mouse platelets demonstrated reduced C-terminal tyrosine phosphorylation of Lyn and Src, further supporting CD148’s role in positively regulating platelet activation.^22^ Taken together, our findings suggest that CD148 enhances platelet activation not only by dephosphorylating PECAM-1 ITIM tyrosines, but also the inhibitory tyrosines of Src and Lyn. These synergistic signaling events combine to amplify GPVI-mediated platelet activation.

This study focused exclusively on the activation of human platelets, however CD148 and PECAM-1 are also co-expressed on other cell types, including B cells, raising the possibility that taFv 179 could also be used to potentiate B cell activation. Exploring the broader applicability of taFv 179 in other blood and hematopoietic cells could provide valuable insights into its potential as a therapeutic tool for modulating immune responses. Future studies examining the effects of taFv 179 and its second- and third-generation products may be an exciting avenue for future investigation.

## Acknowledgements

This work was supported by NIH grant R35 HL-139937

## Authorship Contributions

A.J.M. designed and performed experiments, analyzed data, made the figures and wrote the manuscript. P.J.N. designed experiments, wrote the manuscript, and supervised the work.

